# Parental age selection in *C. elegans* influences progeny stress resistance capacity

**DOI:** 10.1101/2025.04.30.651556

**Authors:** Bennett T. Van Camp, Sean P. Curran

**Affiliations:** Leonard Davis School of Gerontology, University of Southern California

**Keywords:** *C. elegans*, aging, reproductive span, healthspan, stress resistance

## Abstract

With parental age rising around the globe, an increased understanding of the impact on health and longevity is needed. Here, we report how the continuous selection of the last progeny during the *Caenorhabditis elegans* reproductive span results in a diminishment of multiple age-related health measures. After more than fifty generations of late selection, progeny displayed diminished resistance to acute oxidative stress, disrupted partitioning of stored lipids, reduced movement capacity, and an overall shortening of lifespan. In contrast, starvation resistance was improved and late selection had negligible effects on developmental timing and total reproductive output that suggests a reduction in lifespan health to preserve reproductive capacity. The phenotypes of late selection are reminiscent of animals with activation of the cytoprotective transcription factor SKN-1 but are unlikely a result of a spontaneous genetic mutation. These findings suggest the existence of a homeostatic mechanism for bookmarking the temporal boundaries of the parental reproductive span that reshapes the way we think about parental age influencing offspring fitness.

## INTRODUCTION

The underlying mechanisms of aging have been a topic of intense focus by the research community for their impact on multiple issues of societal importance (e.g., health, lifespan, population dynamics, and evolutionary history [1-4]. However, the primary focus of aging research tends to examine the deterioration of a monogenerational cohort of research subjects despite the knowledge that transgenerational effects can be potent drivers of natural aging [4]. Reproductive success is an important life history trait, most often considered in the context of parental fitness and behavior [5, 6]. However, age-related reproductive decline also influences offspring health and fitness [3, 7-9] and parental age has been shown to impact a wide variety of offspring phenotypes, including reproductive output, environmental stress resistance, early life survivability, and lifespan [3, 7-12]. In contrast, over the past fifty-five years, human lifespan has been on the rise in many countries, with a recent decrease observed in the US [13] and a global decrease due to the COVID-19 pandemic [13, 14]. Alongside this trend, the mean parental age has also risen, with the percentage of people becoming first-time fathers over forty doubling [15]. It is imperative to understand the effects of late reproduction have on offspring health but also how these population trends could have on society.

These effects have been observed across taxa, ranging from humans, elephants, roundworms, and flies [3, 7-12]. Several potential mechanisms have been described that may contribute to these effects, including a decline in overall gamete quality, a loss of parental capacity to provide somatic support to eggs and sperm [16], increased mutational load in the germline [17-21], and decreased nutrient provisioning to eggs, as observed in older *Eupelmus vuilleti* mothers [22], or in the case of *Sancassania berlesei*, young mothers lay many eggs in nutrient-rich environments but swap in old age to producing fewer, but larger progeny that are able to compete with their older siblings in a more competitive nutrient environment [23]. However, there is some evidence that this accelerated growth could be detrimental to the younger offspring in the long term [24, 25].

Although the mechanisms underlying the impact parental age have on offspring fitness have been hypothesized, the effects of multi-generational selection of advanced age requires additional exploration. As stated previously, there is some evidence in *Drosophila* that poor genetic quality as a result of advanced parental age could lead to a snowball effect in future generations [19, 20, 26]. However, other studies have shown that selecting for older parental age for several generations in *Drosophila* produces long-lived offspring [27]. The mechanism for these effects, their accumulation, and their durability remain poorly understood, as there appears to be a high variability in the observed effects based on the age of selection mating behaviors [16]. Additionally, in shorter selections (under 10 generations) significant variability has been observed based on the duration of the selection, with the animals initially showing a decrease in reproduction followed by a normalization to baseline [11]. Collectively, we need additional models to better understand how parental age selection impacts health.

To this end, we performed a long-term age selection using *Caenorhabditis elegans* as a model. In order to avoid potential short-term effects and short-term intergenerational variability, we performed this selection for fifty generations before collecting data. Below, we detail the healthspan and lifespan effects resulting from this generational selection, as well as a transcriptomic analyses of the selection cohorts. Here we take a closer look at the ramifications of long-term parental age selection providing a new experimental model and key insights connecting multigenerational selection of late parenthood on offspring health.

## RESULTS

### Repetitive selection of late progeny reduces *C. elegans* lifespan

Advanced parental age is known to negatively impact a broad spectrum of healthspan metrics in offspring fitness(e.g., stress resistance, reproductive output, early life health/survival)[3, 7, 8, 16, 26, 28-32], however, much less is understood about the impact of advanced parental age in a multigenerational context [27, 33]. In order to further study this phenomenon, a late selection paradigm was applied to a population of wildtype (WT) *C. elegans* (**Figure 1A**). In this population (hereafter referred to as WT Late), only progeny produced during Day 3 of parental adulthood were collected and used to create the next generation, representing the last quartile of reproductive output (**Figure 1A**). To assess the impact of repetitive and chronic selection of late progeny, the process was repeated for fifty generations (Gen) “WT Late(Gen50+)” before the changes in physiological and molecular markers were assessed.

**Figure 1:**
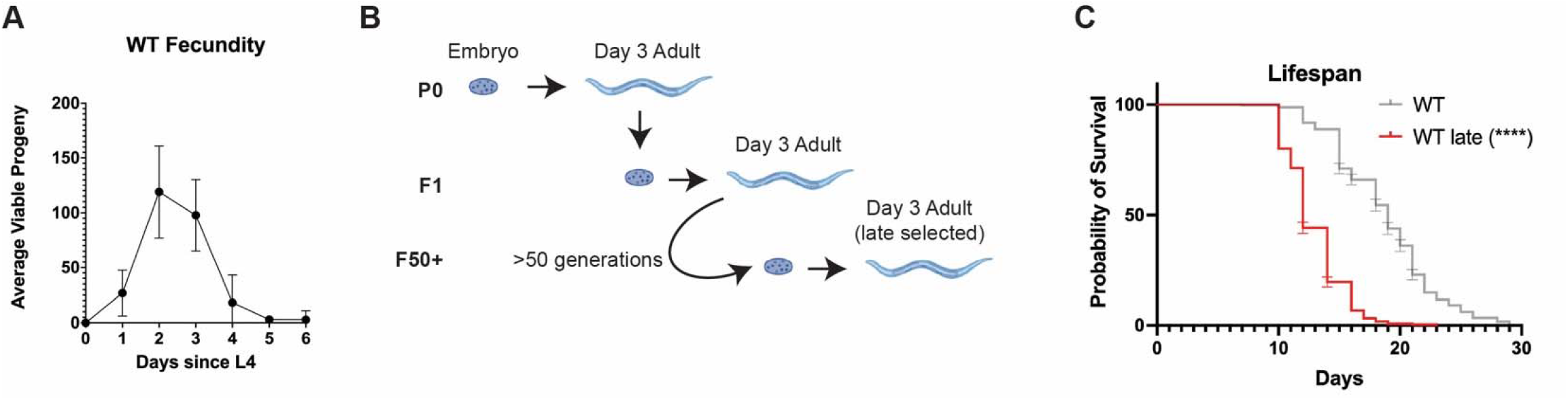
Chronic selection of late progeny shortens lifespan. (**a**) Reproductive curve of WT animals (**b**) Model of the selection process (**c**) WT Late worms exhibit a reduced lifespan compared to WT worms

We first measured the lifespan of synchronized populations of WT Late (Gen50+) animals as compared to unselected controls and discovered that WT Late (Gen50+) worms display a significant diminished life expectancy (**Figure 1C** and Table S1). In fact, the average lifespan of the WT Late (Gen50+) population was 36.84% reduced compared to unselected controls and with a significant reduction of 31.25%, 36.84%, 33.33%, 20.69% at quartile 1, 2, 3, 4, respectively. These data reveal that generational selection of the latest progeny in *C. elegans* drives accelerated aging.

### Repetitive selection of late progeny impacts *C. elegans* healthspan

To understand how the multigenerational selection of the last progeny influences healthspan, we subjected WT Late (Gen50+) worms in a battery of physiological assays to define health status and the relationship, if any, to the decrease in lifespan observed. Movement (e.g., speed) is a powerful surrogate assay for organismal health that declines with age [34-37]. As such, we measured movement capacity of age match WT Late (Gen50+) animals as compared to unselected WT controls. It should be noted that WT late worms are slightly smaller than their WT counterparts, but controlling for this did not impact any of the movement comparisons **(Figure 2A)**. WT Late (Gen50+) worms performed significantly worse than their WT counterparts; displaying a significant reduction in crawl speed and swim speed with no reduction in wave initiation rate and dynamic amplitude. (**Figure 2B-E**). In light of the reproductive selection performed, we also measured the reproductive output of WT Late (Gen50+) worms. Surprisingly, we did not observe a significant change in the total number of progenies in WT Late (Gen50+) worms as compared to WT animals (**Figure 2F**). Although we did note a modest (<5%) delay in the timing of first egg being laid (Figure S1) the general timing of progeny production over the reproductive span was unremarkable (**Figure 2G**). This suggests that at least one aspect of fitness, specifically reproduction capacity, is preserved despite significant changes in organismal lifespan.

**Figure 2:**
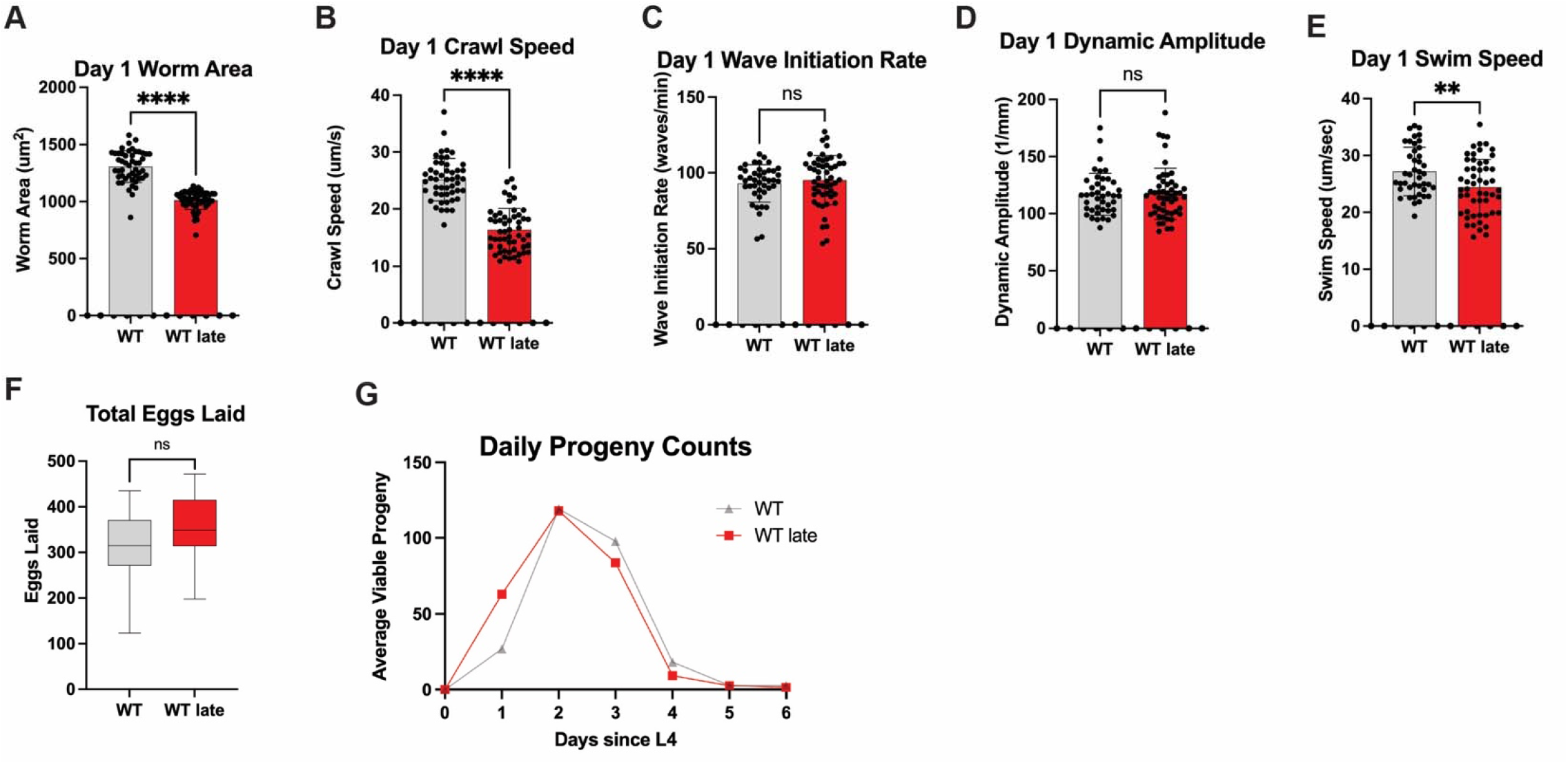
Progeny of late selection maintain reproductive output. (**a**) WT Late worms have less area than WT worms (**b**) WT Late worms have lower crawl speed (**c**) WT Late worms have no change in wave initiation Rate (**d**) WT Late worms have no change in swimming dynamic amplitude (**e**) WT Late worms have lower swim speed (**f**) WT Late worms have no change in reproductive output (**g**) Daily reproductive output of WT Late worms

### Generational selection of late progeny alters oxidative and metabolic resistance capacity

A common hallmark of aging [4] is the impairment of stress resistance capacity and dysregulation of metabolic homeostasis. To further elucidate the impact of long-term late selection on *C. elegans* healthspan, the worms were subjected to oxidative stress via hydrogen peroxide exposure as previously described [38]. At Day 1 of adulthood, WT Late worms performed the same as unselected WT worms (**Figure 3A**). However, when Day 3 adult worms were exposed, WT Late worms performed significantly worse than their unselected counterparts (**Figure 3B**). This data reveals that repetitive selection of the last progeny leads to a loss of the ability to tolerate acute stress with age.

**Figure 3:**
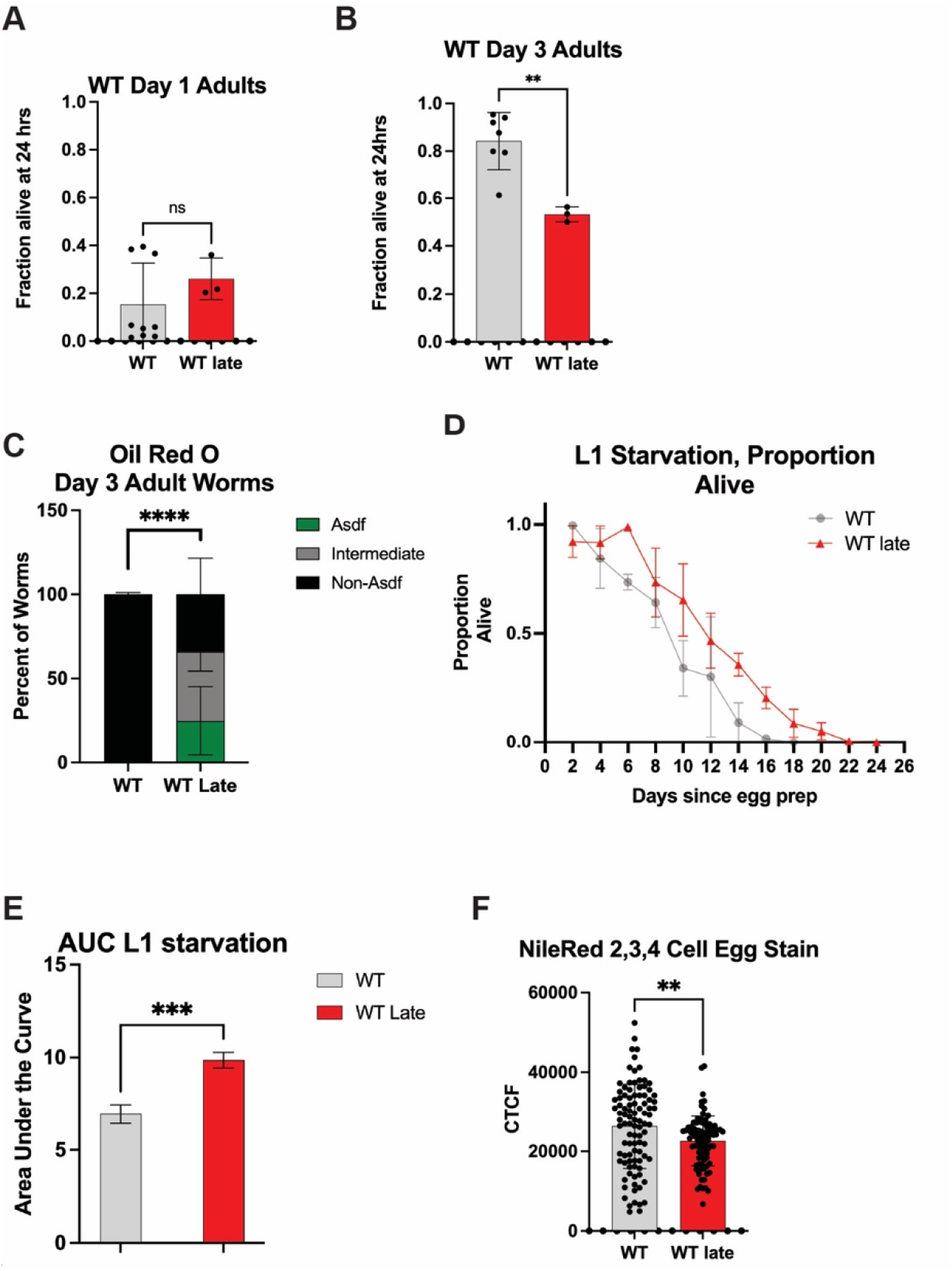
Progeny of late parental selection display altered stress adaptation capacity. (**a**) Day 1 oxidative stress assays show no significant difference between WT and WT Late worms (**b**) WT Late worms have a decreased Day 3 oxidative stress resistance (**c**) Oil Red O staining of Day 3 worms shows an increase in either full or intermediate Asdf in WT Late worms (**d**) L1 starvation assay depicts an immediate decrease in WT survival whereas WT Late only falls off after 6 days (**e**) Area under the Curve of the L1 starvation data (**f**) Nile Red staining shows that WT Late worms have a lower average lipid content

We next assessed lipid distribution across *C. elegans* fat storage tissues by staining with the neutral lipid dye Oil Red O (ORO) [39]; looking specifically for the age-dependent somatic depletion of fat (Asdf), which has been previously described as an intersection between lipid homeostasis and immune response and predictive of lifespan potential [39-41]. Aligned with our previous observation documenting the intriguing relationship with lipid homeostasis and stress resistance capacity we observed a significant decrease in the number of worms that do not display the Asdf phenotype in the WT Late (Gen50+) population as compared to unselected controls (**Figure 3C** and Figure S2A-D); this observation was driven most significantly in worms exhibiting an intermediate phenotype (an incomplete depletion of somatic lipids) (Figure S2C). One hypothesis for why somatic lipids are redistributed to the germline is to ensure the survival of the next generation by providing additional resources. To test this idea, we stress newly hatched larval stage 1 (L1) animals to starvation and measures survival as assessed by the ability to resume development when food is reintroduced [42]. Progeny from WT Late (Gen50+) animals that are derived from germ cells with extra lipid availability, exhibit a significant increase in their resistance to L1 starvation (**Figure 3D,E**). Specifically, unselected WT worms quickly lose the ability to tolerate starvation and resume development when food is reintroduced while WT Late (Gen50+) animals are able to survive the complete absence of food for six additional days without a significant decrease in survival. Although developing oocytes have increased lipid bioavailability, we could not detect a significant increase in total lipids in newly fertilized embryos as measured by quantifying Nile Red stained lipids by fluorescent microscopy. In fact, quantification of two, three, and four cell embryos reveals that WT Late (Gen50+) animals harbor less lipids than unselected WT counterparts (**Figure 3F**) [39]. Additionally, no significant change was measureable in the distribution of lipids between the somatic AB cell precursor or the germ cell precursor P1 cell (Figure S2E,F).

### Repetitive selection of late progeny modifies the steady-state transcriptional landscape

Several of the phenotypes observed in WT Late (Gen50+) animals resemble responses influenced by SKN-1 transcriptional activation [35]. To investigate whether WT Late (Gen50+) animals accumulated a genetic variation that activates SKN-1 we performed genome-wide sequencing (GWS). We identified 78 variants in coding regions, none in known regulators of SKN-1 activity. Nevertheless, we performed RNAi targeting these genes in a strain harboring a *gst-4p::gfp* reporter that is sensitive to SKN-1 activity, but none induced GFP expression (Table S2). Collectively, these data suggest SKN-1 is not activated to the level observed in response to other classical genetic mutations [35].

Despite our inability to clearly define a SKN-1-activated state, in order to better determine the molecular basis of the phenotypes observed <u>in</u> WT Late (Gen50+) animals we next examined the transcriptional landscape by RNAseq. We compared <u>in</u> WT Late (Gen50+) animals to age-matched unselected WT animals which revealed significant transcriptional remodeling in age-matched L4 stage animals (**Figure 4A-C**); 199 with increased expression and 93 with decreased expression. In both the KEGG pathway and the GO term analyses, lipid homeostasis and innate immune response were among the classes of genes that display the most significant change after late selection (**Figure 4B,C**). Specifically, genes involved in innate immune response (e.g., *acdh-1*), lipid transport (e.g., *vit-1, vit-3*), and lipid utilization (e.g., *acs-2*) were affected in late selected progeny (**Figure 4A**). Genes involved in oxidative stress response (e.g., *msra-1*) were also found to be affected in late selected progeny. (**Figure 4A**). Although SKN-1 was not demonstrably activated, transcription factor enrichment analysis (TFEA) revealed targets of SEX-1 and ELT-1 were commonly impacted in late selected progeny (**Figure 4D**) and may contribute to the modified transcriptional landscape that resembles the phenotypes stemming from a modest increase in SKN-1 cytoprotection.

**Figure 4:**
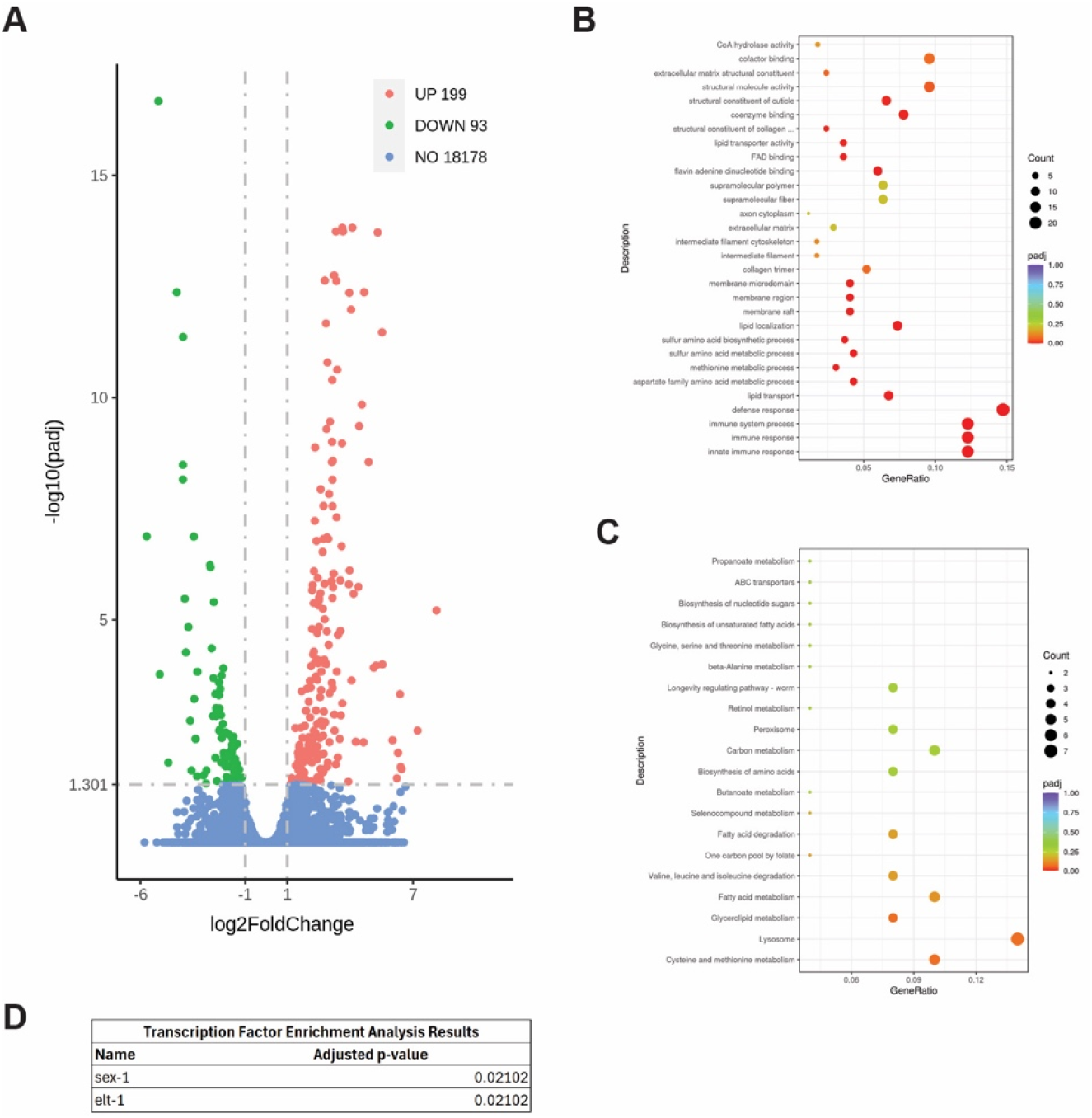
Late progeny selection alters the transcriptional landscape. (**a**) DEG analysis reveals significant changes in both immune response and lipid homeostasis. (**b**) GO term analysis reveals significant changes in immune response and lipid homeostasis (**c**) KEGG pathway analysis reveals significant changes in lipid homeostasis (**d**) Transcription Factor Enrichment Analysis reveals significant changes in the targets of SEX-1 and ELT-1.

## DISCUSSION

Reproductive aging is typically discussed in terms of the health of the parent [5, 6]. However, it has been shown that parental age can also significantly alter the healthspan and lifespan of their progeny [3, 7, 8, 16, 26, 28-32]. We characterized the impacts of a multigenerational change in parental age by 50 generations of selecting for Day 3 progeny, that result in significant alterations to the transcriptomic landscape, diminished overall healthspan, and reduced lifespan.

One of the major changes notices is a vast transcriptomic remodeling of lipid homeostasis genes (**Figure 4A-C**). Lipid changes were some of the most substantial changes in the GO term and KEGG pathway analysis as well as the downregulation of *vit-1* and *vit-3* and an upregulation of *acs-2* (**Figure 4B,C**). This is consistent with the finding that WT Late worms have a lower egg lipid content that their WT counterparts (**Figure 3F**) as a knockdown of *acs-2* has been shown to increase fatty deposits [43] and vitellogenins are well known yolk protein precursors [44]. This is consistent with the hypothesis that long-term parental age effects could occur via decreased nutrient provisioning [16], as has been seen previously in *Eupelmus vuilleti* [22]. However, this result was somewhat surprising given that WT Late worms have better L1 starvation resistance and adult WT Late worms Asdf more, a phenotype usually thought to be a form of terminal investment in the progeny [40] (**Figure 3C-E**). One potential explanation is that while WT Late eggs may not have more lipids, they may be supplied an increased amount of other nutrients from the parents. Another potential explanation is that WT Late worms develop slightly slower and could use up their lipid stores slower that WT Late worms (Figure S1A). Separately, Transcription Factor Enrichment Analysis reveals that WT Late worms displayed an enrichment for the targets of SEX-1 and ELT-1 respectively. This is interesting because SEX-1, nuclear hormone receptor, is known to impact early development [45] and ELT-1 is an erythrocytelike transcription factor which is known to impact the expression of SKN-1 targets [46, 47]. It is possible that the affected targets of these transcription factors are at least partially responsible for the reproductive and SKN-1-related phenotypes of WT Late worms. However, none of the GWS hits individually was sufficient to drive a change in *gst-4::gfp* expression when knocked down via RNAi (Table S2).

The WT Late population also exhibited several other changes, including a decrease in late life stress resistance, lower Day 1 crawl speed and swim speed, and a 36% decrease in median lifespan (**Figure 3;** Table S1). This was surprising as similar studies in *Drosophila* found that long-term parental age selection improved lifespan and stress resistance [27]. One potential explanation is that the severity of the selection has a significant impact on the outcome of the selection. Here, the selection was much less severe, beginning just after the drop from peak reproduction while previous studies use a much more extreme cutoff [27]. Another note of interest is that this decreased lifespan, healthspan, and stress resistance phenotypes all mimic patterns that have been previously observed in mutants with a constitutively active stress resistance pathway, perhaps indicating that the cumulative effect of late parental age selection is somehow stressful to the animals [35, 40, 41]. We selected for a minimum of fifty generations to minimize possible epigenetic effects that could confound analyses that typically last 3-5 generations [48, 49]. Although our genome-wide sequencing identified possible mutations, unsurprisingly none were homozygous, which suggests that the changes observed are not a result of genetic selection. Collectively, this study reveals the repetitive and chronic selection of late progeny across generations can significantly alter healthspan and lifespan trajectories.

## MATERIALS AND METHODS

### Maintenance of *C. elegans* Strains

Strains were grown at 20° C on nematode growth media (NGM)+streptomycin plates with OP50 food. WT, N2 Bristol strain was used and is the strain WT Late was derived from. All strains were unstarved for at least three generations before use.

### RNAseq Analysis

Worms were synchronized overnight as L1s and then dropped on NGM+streptomycin plates with OP50 for 48hrs and then collected. They were then washed 3x with M9 buffer and frozen at -80°C in TRI reagent until use. Worms were then homogenized and had their RNA extracted using the Zymo Direct-zol RNA Miniprep Kit (Cat. #R2052). Samples were then sequenced and read counts, Differential expression analysis, GO term analysis, and KEGG pathway analysis were reported by Novogene.

### Oil Red O Staining

Staining was performed as previously described [35, 39]. In brief, worms were synchronized overnight as L1s and then dropped on NGM+streptomycin plates with OP50 for 120hrs and then collected. They were then washed with PBS+triton, then rocked in 40% isopropyl alcohol for 3 mins. Worms were then pelleted and treated with ORO in H2O for 2 hours. They were then washed in PBS+triton for 30 minutes and imaged at 20× using LAS X software and Leica Thunder Imager Flexacam C3 color camera. The stained worms were categorized as Asdf (complete somatic lipid depletion), intermediate (incomplete somatic lipid depletion), and non-Asdf (no somatic lipid depletion).

### Starvation Resistance Assay

Worms were egg prepped and synchronized overnight as L1s and then rotated slowly in M9 at 20°C. 10uL were removed every 2 days and the larvae bodies were counted on a NGM+streptomycin plate with OP50. They were then allowed to grow for 72 hrs and recounted to get the proportion alive. The experiment ended once a 0% survival was reached for all conditions.

### Egg Nile Red Staining

Staining was performed as previously described [39]. Worms were egg prepped and then the eggs were washed in PBS+triton. Eggs were then incubated in 40% isopropyl alcohol overnight. The next day, the eggs were moved into Nile Red+DAPI staining solution and incubated for 2 hours in the dark. They were then washed with PBS+triton and imaged at 63X using LAS X software and Leica Thunder Imager. Corrected total cell fluorescence (CTCF) was then measured in ImageJ and Microsoft Excel.

### Oxidative Stress Assay

Assay was performed as previously described [35, 41]. Synchronous populations of Day 3 adults were washed 3x in M9+triton. The worm pellet was aspirated down to 500ul and then 500uL of 20mM hydrogen peroxide was then added. They were then incubated on a rotator at 20°C for 25 minutes. Worms were then washed 3x with M9+triton and dropped onto NGM+streptomycin plates with OP50. Worms were then counted and then counted again 24 hrs later to get their survival.

### Movement Measurements

For all assays, worms were egg prepped and synchronized overnight as L1s. Worms were then added to an OP50 plate and allowed to grow for and 72hrs to Day 1 adults. Worms were then washed with M9+triton onto an unseeded NGM plate. The worms were allowed to acclimate for 30+ minutes to allow the liquid to evaporate. Crawling videos were then taken for 1 minute at 7.5 fps. All imaging was done with the MBF Bioscience WormLab microscope. Analysis was then performed by WormRACER [37].

### Lifespan Assay

Assay was performed as previously described [36, 50, 51]. Worms were egg prepped and synchronized as L1s overnight. They were then dropped onto NGM+streptomycin and OP50 plates. Worms were moved periodically to remove progeny as needed and kept at 20°C. Worms were scored via prodding with a platinum wire daily. Bagging, vulval bursting, and desiccation on the side of the plate led to censorship of that worm.

### RNAi Assay

*gst4::gfp* reporter worms were egg prepped and synchronized in M9 overnight at 20°C. L1s were then dropped onto NGM plates seeded with the appropriate L4440-based RNAi clone. At L4 stage, worms were assessed for changes in reporter expression and scored on a three-point scale (0 OFF, 1 normal, 2 increased, 3 HIGH).

### Developmental Timing Assay

Worms were egg prepped and synchronized in M9 overnight at 20°C. Single L1s were then put onto NGM+streptomycin plates with OP50. Worms were then scored every 2 hours until they lay their first egg.

### Reproductive Output Assay

Worms were egg prepped and synchronized in M9 overnight at 20°C. L1s were then put onto NGM+streptomycin plates with OP50. L4s were then singled. Each day, eggs were counted, and the adult worm was moved to a new plate. Assay continued until egg laying ceased.

### Statistical Analysis

All statistical analysis was done on GraphPad Prism version 9.5.0. Statistical analysis done via Prism were either an unpaired t-test or a Kaplan-Meier simple survival analysis. p<.05 was the threshold for significance.

## ACKNOWLEDGMENTS

We thank S. Keel and T. Phan for technical assistance. This work was funded by the NIH R01AG058610 and Hevolution Foundation award HF AGE-004 to SPC and NIH T32AG052374 to BTVC. Some strains were provided by the CGC, which is funded by the NIH Office of Research Infrastructure Programs (P40 OD010440). We thank WormBase for database curation and data access.

## Author contributions

Conceptualization: BTVC and SPC; Methodology: BTVC and SPC; Investigation: BTVC; Visualization: BTVC and SPC; Supervision: SPC; Writing (original draft): SPC; Writing (reviewing & editing): BTVC and SPC.

## Data and materials availability

All data are available in the main text or the supplementary materials.

## SUPPLEMENTAL FIGURES

**Figure S1:**
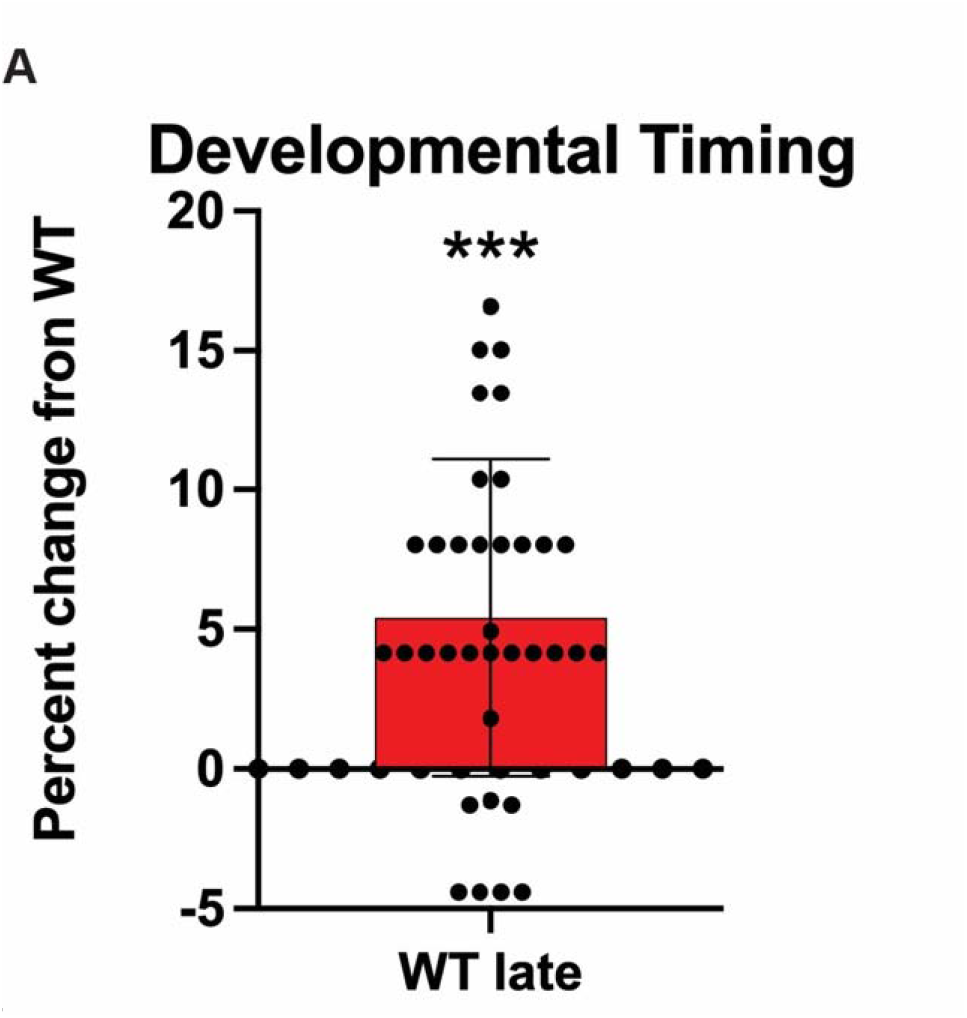
(**a**) WT Late worms have a slightly increased time till first egg lay compared to WT worms

**Figure S2:**
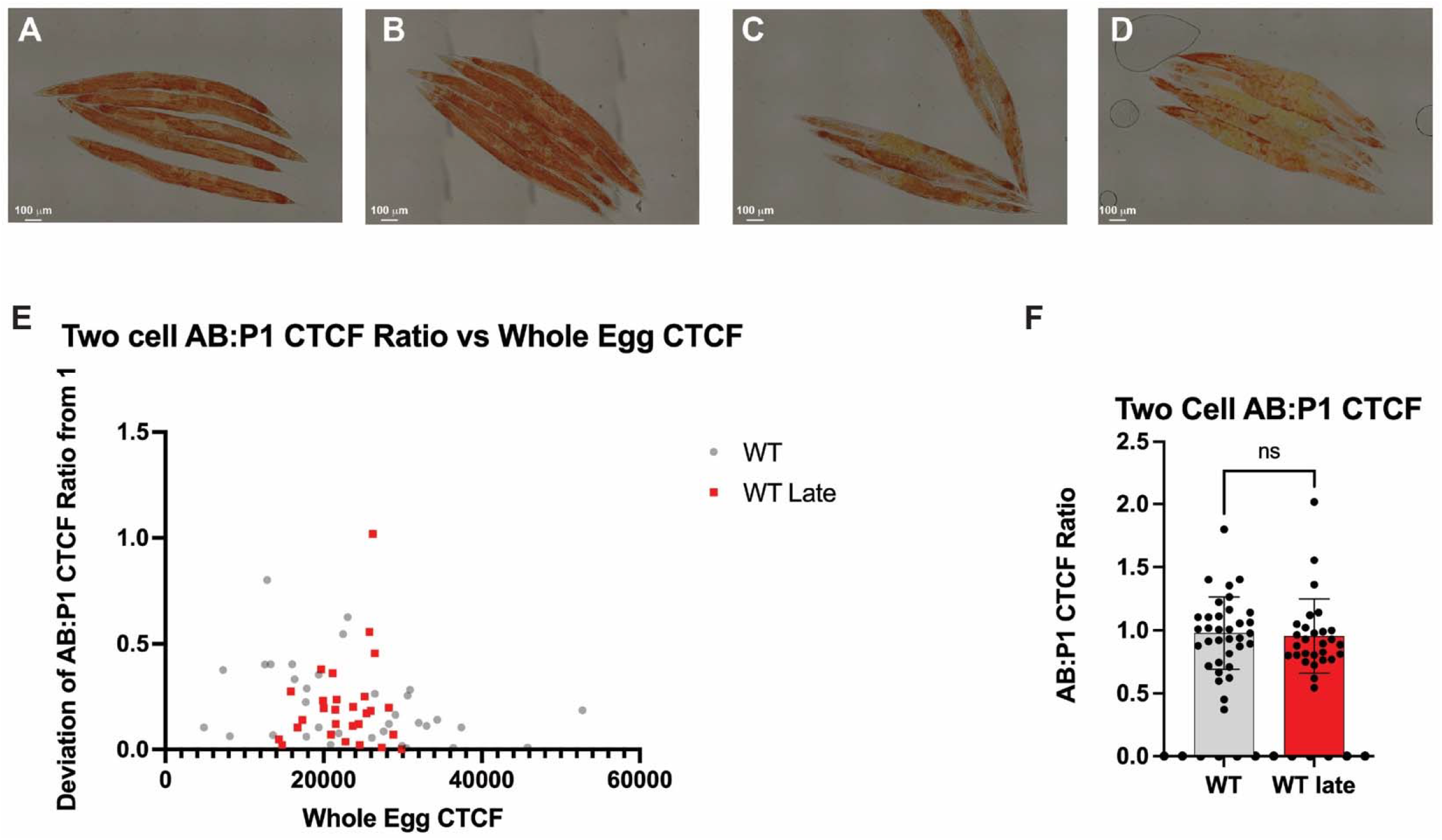
(**a**) WT non-asdf worms (**b**) WT Late non-asdf worms (**c**) WT Late intermediate Asdf worms (**d**) WT Late Asdf worms (**e**) WT Late worms have no significant change in the distribution of Nile Red stained lipid distribution between their AB and P_1_ cells and the distribution is not dependent on the total lipid content of the egg (**f**) WT Late worms have no significant change in the distribution of Nile Red stained lipid distribution between their AB and P_1_ cells

## SUPPLEMENTARY TABLES

<u>Table S1:</u> Quartiles for WT and WT Late lifespan assay

<u>Table S2:</u> *gst-4::gfp* reporter RNAi of GWS hits results

