## Supplementary figures and images for "Parental age selection in *C. elegans* influences progeny stress resistance capacity"

### Table S1

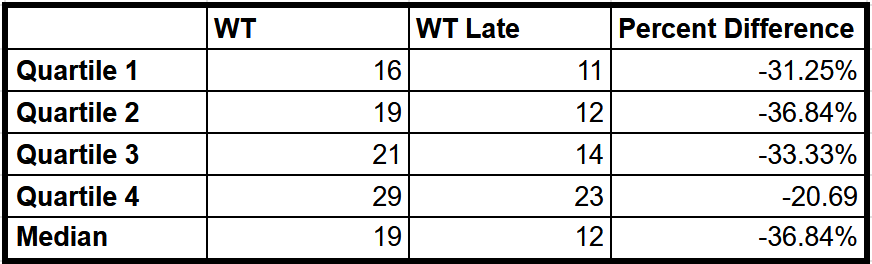
